# Computational Design Map for Heterogeneous Experimental Studies

**DOI:** 10.1101/2021.05.25.445627

**Authors:** Chhaya Kulkarni, Nuzhat Maisha, Leasha J Schaub, Jacob Glaser, Erin Lavik, Vandana P. Janeja

## Abstract

This paper focuses on the discovery of a computational design map of disparate heterogeneous outcomes from bioinformatics experiments in pig (porcine) studies to help identify key variables impacting the experiment outcomes. Specifically we aim to connect discoveries from disparate laboratory experimentation in the area of trauma, blood loss and blood clotting using data science methods in a collaborative ensemble setting. Trauma related grave injuries cause exsanguination and death, constituting up to 50% of deaths especially in the armed forces. Restricting blood loss in such scenarios usually requires the presence of first responders, which is not feasible in certain cases. Moreover, a traumatic event may lead to a cytokine storm, reflected in the cytokine variables. Hemostatic nanoparticles have been developed to tackle these kinds of situations of trauma and blood loss. This paper highlights a collaborative effort of using data science methods in evaluating the outcomes from a lab study to further understand the efficacy of the nanoparticles. An intravenous administration of hemostatic nanoparticles was executed in pigs that had to undergo hemorrhagic shock and blood loss and other immune response variables, cytokine response variables are measured. Thus, through various hemostatic nanoparticles used in the intervention, multiple data outcomes are produced and it becomes critical to understand which nanoparticles are critical and what variables are key to study further variations in the lab. We propose a collaborative data mining framework which combines the results from multiple data mining methods to discover impactful features. We used frequent patterns observed in the data from these experiments. We further validate the connections between these frequent rules by comparing the results with decision trees and feature ranking. Both the frequent patterns and the decision trees help us identify the critical variables that stand out in the lab studies and need further validation and follow up in future studies. The outcomes from the data mining methods help produce a computational design map of the experimental results. Our preliminary results from such a computational design map provided insights in determining which features can help in designing the most effective hemostatic nanoparticles.

## I. INTRODUCTION

In this study, we focus on discovering a computational design map from multiple disparate lab experiments from the porcine experiment data collected after hemostatic nanoparticles are injected onto the trauma-induced porcine animals. Our study focused on levels of immune response variables that impacted the survivability factor along with clot formation in the porcine subjects as a result of administration of different hemostatic nanoparticles. The aim is to identify hemostatic nanoparticles that result in blood clotting and survival of the animal as a function of the variables observed for a particular hemostatic nanoparticle. The data from experiments is small but at the same time fairly complex and needs careful pre-processing to reflect the biological processes at play. For example, instead of the biological variable observed directly, it may be more important to use the change in the variable over time as a factor in the data evaluation. This required careful collaboration among the data science and bioinformatics scientists. Our work is motivated by a domain which may have far reaching impacts to designing clinically translatable hemostatic nanoparticles while maintaining its safety and efficacy. This understanding can help with situations of blood loss in combat scenarios and also in devising a data driven framework to study immune responses during a cytokine storm resulting from a trauma event.

### Motivating Scenario

*Assessment of blood loss in trauma related injuries is of prime importance. Combat situations can result in internal bleeding and there may be lack of definitive care (medical resources). Not just on the battlefields alone but many blunt traumas including care accidents can result in internal bleeding in humans. Administration of hemostatic nanoparticles has been found to promote faster blood coagulation at injury sites thereby saving lives. However, even when coagulation is accelerated, if other factors like inflammation occur, bleeding may still continue leading to lethal outcomes. Thus understanding the relationships between the nanoparticles and the outcome variables plays a vital role as it can help in the rapid development of nanoparticles which can prove to be effective in trauma situations.*

### Study setup

*Experiment in the lab setup involved intravenous administration of hemostatic nanoparticles or controls in 19 porcine animals. The reason behind choosing the porcine models for such studies is mainly the anatomic and histopathologic similarities with humans, and as porcine animals show activation for the same dosages as complement reactive humans, porcine animals are a feasible option without provoking hypersensitivity reactions. Data was recorded for 19 porcine animals. Blood samples were collected and weighed at each minute to determine the amount of blood loss* experienced *by the animals. Cytokine analysis along with cellular analysis and clotting analysis were all done. Understanding what levels of cytokines are needed can prove beneficial not only for blood coagulation but also for future applications in understanding potential roots of cytokine storms such as in COVID19. The cytokine storm following sepsis has been proven to be an important mechanism for triggering acute respiratory distress syndrome, which is a fatal uncontrolled systemic inflammation characterized by high concentrations of pro-inflammatory cytokines and chemokines, secreted by immune effector cells. The cytokine storm also occurred in the recently emerged novel coronavirus disease (COVID-19). Therefore, cytokines which usually help the immune system to fight infections are potentially harmful in the course of COVID-19 infections. Studies have highlighted that avoiding or mitigating the cytokine storm may be a key treatment for severe acute respiratory syndrome coronavirus [4,5].*

Our framework moves in this direction by providing a data centric way of analyzing the complex connections between heterogeneous lab experiments and finding the common patterns that can be further investigated in the lab. Several data mining experiments in hematology and other biochemistry based experiments are inclined towards predicting the disease outcome [11,12]. Our study not only uses a predictive model, but it also looks in depth at the features of the in-vitro data individually and in combination to determine which features influence the target variable such as clot formation. Also, majority of the experiments have access to a large sized dataset, whereas, we worked our way through a small but highly heterogeneous real world dataset obtained from a laboratory set up and yet we were able to discover interesting results [12-14], In addition, the way we have preprocessed the data in our study is most certainly different from the traditional preprocessing tasks such as eliminating unimportant attributes, dropping missing values, imputing missing values that are typically followed in other studies [12-14]. We performed our experiment and analysis on porcine data using the popular tools which are widely used in data mining on bioinformatics data [14]. We have incorporated the use of computational design map in our study which is an interesting way to find commonalities among the different experimental results. Implementation of data mining methodologies for knowledge discovery on hemostatic nanoparticles data is an unexplored area. Through our study we hope to bridge this gap.

Our contributions include the following:

- We provide a collaborative framework to connect hetero- geneous lab experiments with different nanoparticles. We form a computational design map connecting the various variables resulting from disparate experiments.
- We use an ensemble approach to combine results from multiple data mining methods to augment our findings and identify variables of the hemostatic nanoparticles experiments that are important.

Our visualization highlights the discovery of variables that are on a critical path between multiple patterns found across disparate experiments.

The rest of the paper is organized as follows. In section 2 we discuss our overall methodology. In section 3 we discuss the experimental results. Finally, we conclude and discuss the future directions for this study.

## II. METHODOLOGY

The key aim of this study is to identify key features from those measured across the various experiments and their relationships to understand the impact of the hemostatic nanoparticles. The experiments were carried out on 19 porcine animals, which were further classified into three groups namely vehicle control, control nanoparticle and hemostatic nanoparti- cle groups, depending on whether they belonged to a control group or recieved the experimental nanoparticle. There were several results measured from all these animals under various stages of trauma recorded in the animals. The variables included measurements from cytokine and non cytokine features. We wanted to evaluate whether a certain presentation in a variable results in survivability of the animal or clotting in the animal. For this we created decision trees to derive rules from the porcine data. Additional comprehensive rules from the porcine data were generated using Apriori algorithm to obtain class based association rules. We also ran our data through a feature ranking algorithm to establish the important features in our porcine data. Association rules and decision trees contributed to the overall computational design map showing the plausible factors by importance and their connection to outcomes of survivability and clot formation in the animals.

Since this data set is relatively small as compared to the traditional data science settings we did not evaluate a Bayesian machine learning model [1,8,9]. In addition, we did not want to start with any independence or dependence relationships but wanted to mainly focus on associations since we did not have prior assumptions about any of the eleven features

The overall methodology is illustrated in Figure 1. Specific sequential actions carried out are discussed next.

**Fig. 1.**
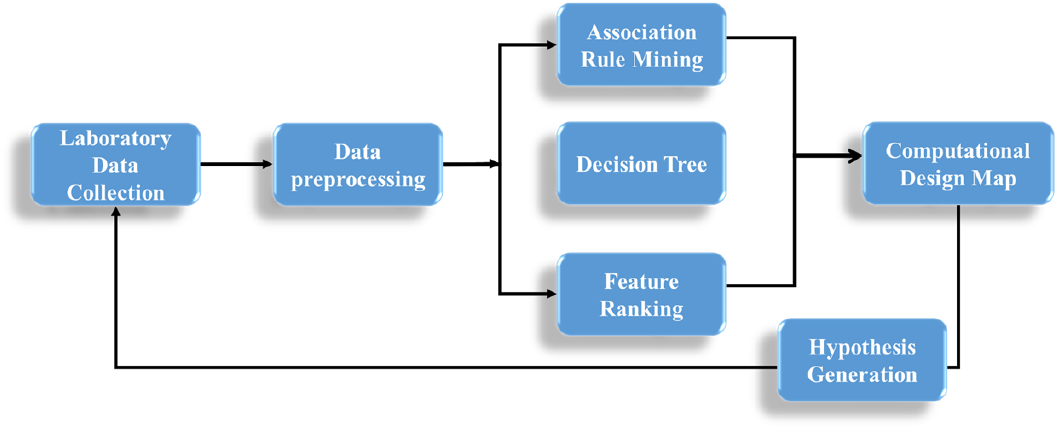
Overall Methodology

### A. Laboratory Data Collection

Our experiment involved intravenous administration of hemostatic nanoparticles in 19 porcine animals and their data was recorded. These 19 animals were divided into 3 groups. The three groups consisted of control, hemostatic and vehicle control groups. Animals were observed after they underwent a trauma incision. Blood was collected from the abdominal cavity continuously and weighed every minute to quantify blood loss. Blood samples were collected every 15 minutes and analyzed for IL6, IL8, Neutrophil, Monocytes, Lymphocytes, TNF-alpha, INF-alpha and IL1-beta. This data was structurally collected. Biochemists also noted which animals survived and which did not; animal’s clot outcome was also recorded.

### B. Data Preprocessing

All the data from the laboratory contained continuous values. This data had to be transformed to perform data mining. We prepared two subsets of data first from the temporal blood loss data, i.e. blood loss for every minute as a time series, second from the cytokine and non-cytokine variables measured every 15 minutes. Our approach here uses this feature data. In these datasets we created two subsets to study changes over time. In one subset we computed the averaged differences of the adjacent time intervals and for the other subset we computed the differences with respect to the baseline value. For each of these datasets, we then converted the obtained averaged values into three ranges of low, medium and high by setting a threshold. We discretized the data to run the decision tree algorithm for classification of the target feature clot outcome or survivability. The preprocessed data is encoded before applying it in certain models such as in the case feature importance model. In association rule mining and in decision tree classification we used the discretized categorical data.

It is important to note that the data that we were working with was skewed. We had a dataset consisting of 19 animals with their corresponding blood loss and cytokine feature data. In the feature data we could observe the imbalance in the data clearly in the target variable’s class representation, we had seven animals with blood clot formation and twelve animals with no blood clot formation. We used an ensemble approach with multiple levels of analysis to evaluate different facets of this data. Our prior work in collaborative ensemble learning has led to promising results in such heterogeneous datasets [15].

### C. Association Rule Mining

Association rules are utilized to understand what relationships are present between the dependent and independent variables in the dataset. The association rule mining algorithm can be broken down into two steps. All frequent itemsets are found in the first step. The frequent item set is the item set that is included in at least minimum support transactions. The association rules with the confidence at least minimum confidence are generated in the second step.

In addition to generating association rules we also generated a visualization (arules,arulesViz [3]) depicting the connection between the various features to generate the association rules on our porcine data. We ran the Apriori algorithm on our processed data to generate the association rules. We set all the independent features as the antecedent and clot outcome as the consequent. This led to the generation of a large number of association rules, where the independent features formed the left hand side of the rules and ‘clot outcome=yes’ was present on the right hand side of the rules. We did similar experiments with ‘survivability’ as an outcome.

### D. Decision Trees

In our study, we applied decision tree algorithms on the porcine data twice, once with clot outcome as the target variable and second with survivability of animals as the target variable. Both these cases led us to ascertain which features of the porcine data play an important role in blood coagulation and survivability of the porcine animals. We used the J48 algorithm to construct the decision trees.

### E. Feature Ranking

Feature importance is a process that involves assigning a score to independent features based on how useful they are at predicting the dependent variable. We used feature ranking algorithms to determine the attributes that were most prominent in the porcine data. We used information gain and ranking for evaluating our features. We set the clot outcome feature as the target variable to determine which features are important for the clot outcome.

We also used feature importance scores, such as statistical correlation scores, coefficients calculated as part of linear models, decision trees, and permutation importance scores [2, 7]. In this experiment, we built a model using ensemble technique namely random forest on the blood loss data of the animals. We derived interesting results using feature importance functionality. We also used the permutation importance of features on our dataset.

### F. Design Map

We utilized the decision trees, association rules and feature ranking to conclude the interconnection between various features and the outcomes. Moreover the feature importance from each method is combined to provide a higher confidence in the features which should be evaluated further in additional lab experiments. We refer to this interconnected set of critical features as computational design maps. This type of a collaborative ensemble approach to generate a computational design map is useful in forming connections and interconnections when there is a heterogeneous collection of data and results available. Design maps provide clarity in identifying commonalities across experiments. We have conducted many collaborative, ensemble experiments in our effort towards finding which factors are necessary for clot formation. In each experiment, we had one or two key attributes emerging across experiments. Forming connections among the features from different experiments was a challenge in traditional methods. Computational design map addressed this problem by helping us locate which of the independent features kept occurring repetitively and which features occurred in combination with other features. Design maps helped us communicate to the scientists which factor seemed most likely in blood clot formation and survivability and our findings have been echoed by existing literature as well showing the potential efficacy of our proposed models. This helped form novel hypotheses to evaluate in the lab experiments.

Input: Temporal Porcine Data(T_p_)

Output: Transformed Data for Machine Learnings Models

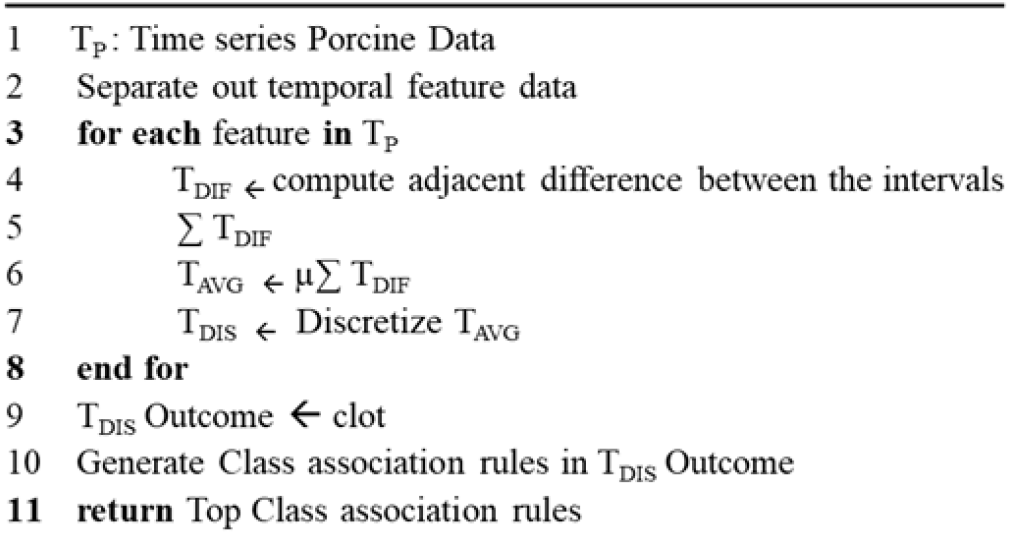

Algorithm 1 shows data preprocessing for the initial temporal feature data. We compute the adjacent difference between the intervals for every feature of the data. We proceed further to calculate the sum of all the adjacent differences between the intervals and then compute the average of these differences. This is used to generate a high, medium, low range for each feature.

We next use the discretized Adjacent Time series Porcine Blood Loss Data denoted as TAV G. We apply Apriori algorithm over TAVm data. We set the values for support, confidence and number of rules we would like our algorithm to derive. We set the clot outcome as the consequent and the remaining data as the antecedent. We then filter out the non-redundant set of association rules and identify the top class association rules.

## IV. EXPERIMENTAL RESULTS

We employed a collaborative ensemble of techniques in order to determine the factors responsible for blood coagulation. All the experiments that we carried out fell in the supervised domain of machine learning and in each experiment we set the target variable - clot outcome or survivability. We next outline our experimental results. The distribution of our data in terms of clot and survival outcomes is shown in table I.

**TABLE I:** Target Variable Distribution in Data

| Target Variable | Yes | No |
| --- | --- | --- |
| Clot Outcome | 7 | 12 |
| Survived | 12 | 5 |

1. **Association Rule Mining:** Figure 2 shows a network containing the top ten association rules when the clot_outcome feature is selected as the consequent. The independent features of the dataset are the vertices and the rules form the edges. We can notice from the figure that all the edges are directed towards the center clot_outcome since we have selected the clot_outcome as the consequent. We observed that a high value of IL6 has been largely responsible for the clot formation. We can also see in one of the rules that most animals that have survived had clot formation. Larger circles denote that they have high support value. All the ten rules depicted in the figure have the same value for lift.
2. **Decision Trees:** Figure 3 illustrates the decision tree generated on porcine data. Average_IL6 is the root node and it further splits down into INF-alpha. Thus, our decision tree algorithm gave us two rules for explaining the clot formation phenomenon which are summarized below:
  1. If Average_IL6=High then Clot formation occurs in animals
  2. If Average_IL6 is not high and INF-alpha is low then clot formation occurs in animals.
  As per decision tree result analysis, IL6 has been the most important factor for blood coagulation.
3. **Ranking Attributes:** Based on the information gain and ranking we extracted the ranked list of features and their scores. Features were listed based on their scores from high to low from top to bottom respectively. Table II shows the attributes with the scores below. In this experiment IL6, Lymphocyte, INF-alpha were the top three ranked attributes.

**Fig. 2.**
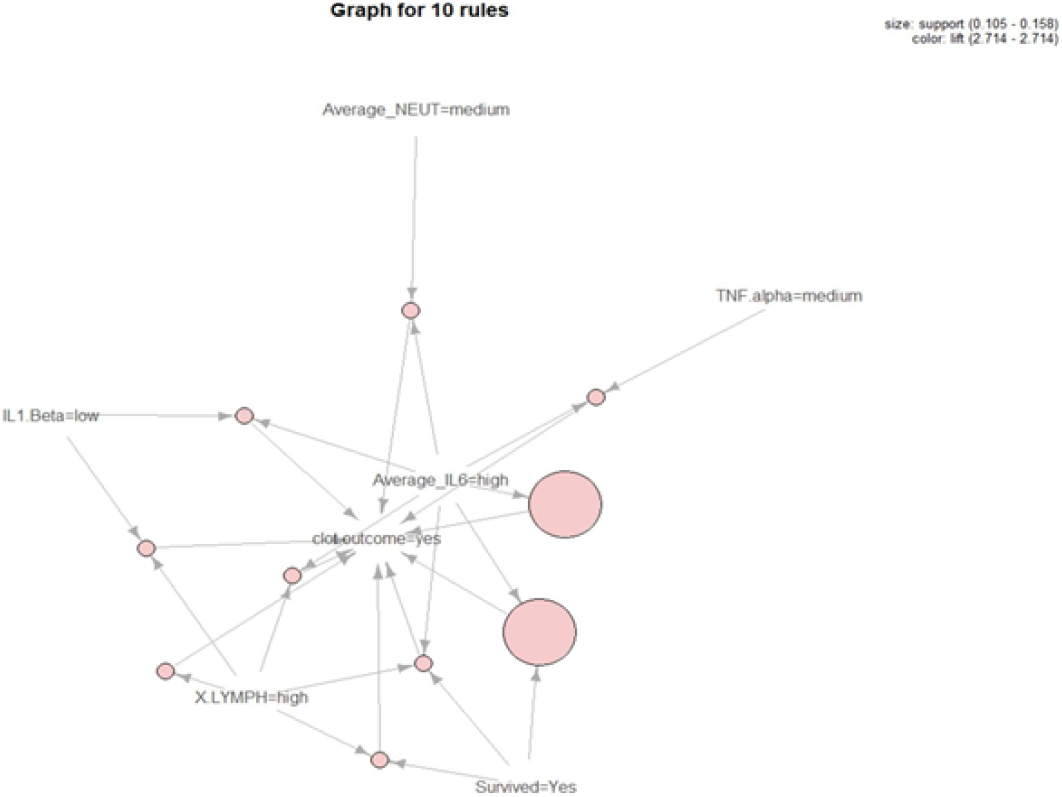
Graph based visualization with items and rules as vertices

**Fig. 3.**
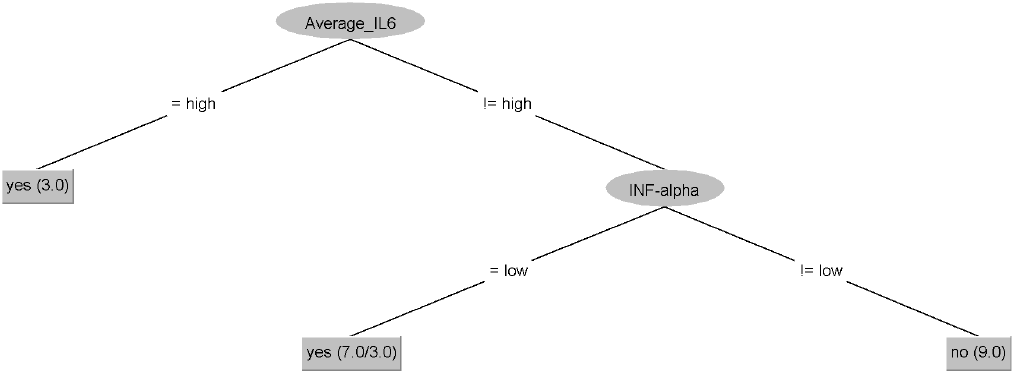
Decision Tree on the Porcine Data

**TABLE II:** Ranked Attributes

| Number | Score | Feature |
| --- | --- | --- |
| 1 | 0.31872 | Average IL6 |
| 2 | 0.19489 | #LYMPH |
| 3 | 0.1588 | INF-alpha |
| 4 | 0.14708 | TNF-alpha |
| 5 | 0.09989 | Average IL8 |
| 6 | 0.07949 | #MONO |
| 7 | 0.07629 | Average NEUT |
| 8 | 0.05796 | Survived |
| 9 | 0.0527 | IL1-Beta |
| 10 | 0.00635 | Average BloodLoss |

In our experiment we also made use of the Random Forest algorithm for feature importance (implemented in Scikit-learn package as RandomForestClassifier classes). After we fit the model, we use the feature importances that can be accessed to retrieve the relative importance scores for each input feature. The figure 4 shows the features along with their scores. We can see that IL6, INF-alpha, Lymphocyte, IL8 and bloodloss are the important features as per the features importance. When we look at the permutation of features importance on porcine data we saw that IL6 was again the most salient feature of the porcine data. Features along with their features importance scores have been shown in figure 4.

**Fig. 4.**
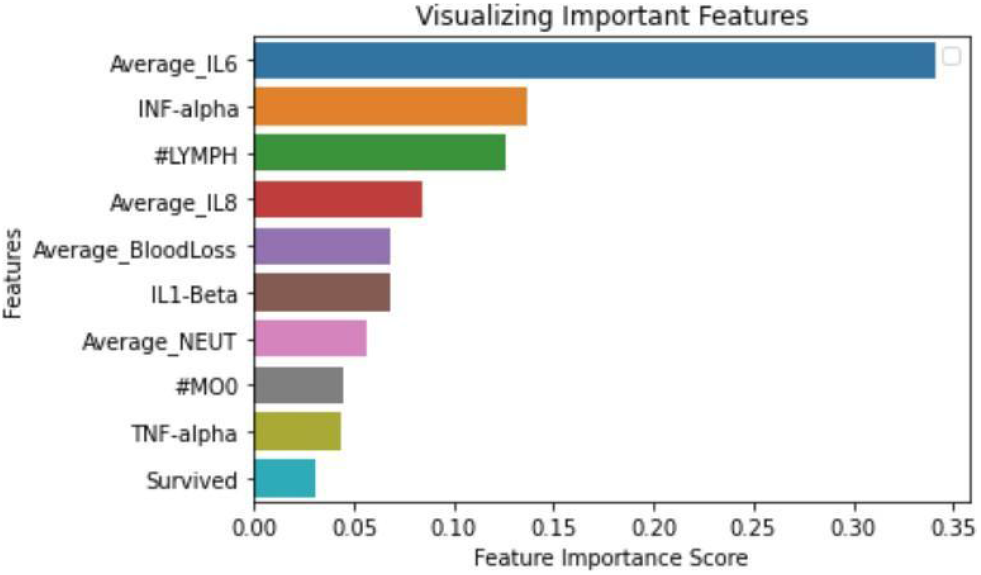
Features Importance applied on Porcine Data

### A. Computational *Design Map*

Our approach for a heterogeneous set of experiments was to consider a collaborative ensemble to identify key features that need to be further evaluated with novel hypotheses.

As depicted in figure 5, we came to discover a computational design map to help us comprehend which attributes are common across the experiments and which attributes of the study are uniquely identified by an experiment.

**Fig. 5.**
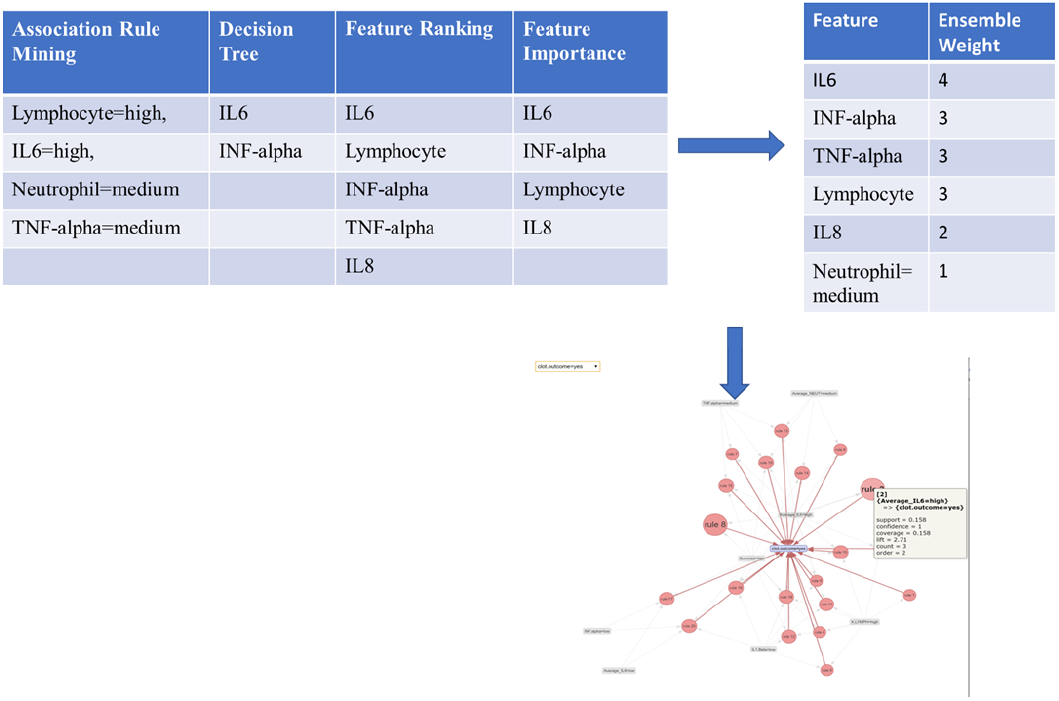
Computational design Map for outcome = Clot

**Fig.6.**
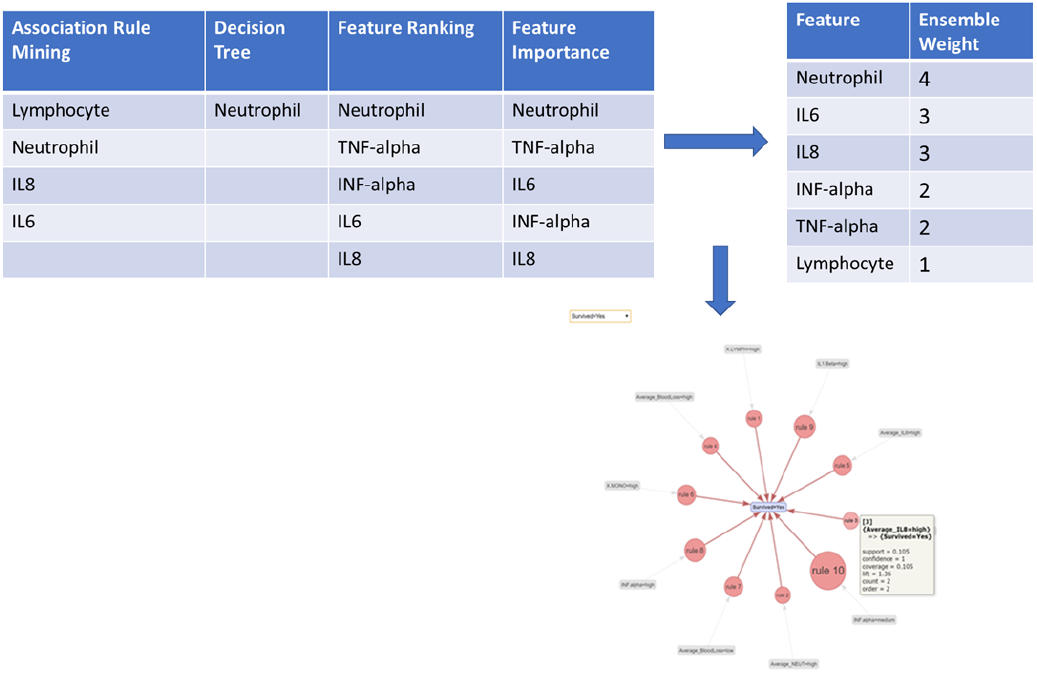
Computational design Map for outcome=Survived

We can observe that IL6 is common across all the individual methods when the outcome was clot, giving it a high weight, whereas Neutrophil has the lowest weight. This results in hypothesis development for hemostasis and inflammation. In the case of the outcome as survived also IL6 gets a high weight. However, Neutrophil also receives a high weight unlike in the clot outcome. Increases in neutrophils are a normal result of trauma, but their presence alone is inconclusive. Their activity in the trauma is critical to outcomes, and the cytokine levels, particularly of IL-6 and IL-8 are essential for understanding whether they are contributing to resolving injury or exacerbating outcomes [16]. The fact that neutrophils are high means that they have not been exhausted.. When neutrophils are exhausted, negative outcomes are often seen in trauma with progression of injury (Same ref as last) Specifically, these findings give us new insight into which factors are critical so we can perform in vitro screening across classes of nanomaterials and determine the interactions between molecular features and negative outcomes associated with infusion reactions.

## V. CONCLUSION & FUTURE WORK

We observed that IL6 was one feature that cropped up in all our experiments. We concluded that a high value of IL6 helps in blood coagulation. Along with IL6, another important feature responsible for blood coagulation was Lymphocyte. When Lymphocyte value was in the high range it triggered blood coagulation. These findings helped identify which features were essential in the blood clot formation and also in the survivability factor. This helps us steer towards the right direction for developing the hemostatic nanoparticles with the right composition. We could also see how effective data science experiments can be to help bioinformaticians to make better decisions. Design maps helped us find the overlapping results from different experiments to generate novel hypotheses for further evaluation. In our future work we will be focused on collecting more data and building a model that predicts the clot formation as well survival of animals based on its hematology report. We also believe that the generated association rules can be used in the future towards building association-rule based predictive models [6].

## V. ACKNOWLEDGEMENTS

Authors Lavik and Maisha would like to acknowledge AIMM Research award (DOD Award Number# W81XWH1820061).

## Notes

### Competing Interest Statement

The authors have declared no competing interest.

